# Peptidoglycan sensing prevents quiescence and promotes quorum-independent growth of uropathogenic *Escherichia coli*

**DOI:** 10.1101/830877

**Authors:** Eric C. DiBiasio, Hilary J. Ranson, James R. Johnson, David C. Rowley, Paul S. Cohen, Jodi L. Camberg

**Affiliations:** Department of Cell & Molecular Biology, The University of Rhode Island, Kingston, RI 02881; Department of Biomedical & Pharmaceutical Sciences, The University of Rhode Island, Kingston, RI 02881; Minneapolis Veterans Health Care System, Minneapolis, Minnesota, USA; Department of Medicine, University of Minnesota, Minneapolis, Minnesota, USA

**Keywords:** quiescence, dormancy, proliferant, peptidoglycan, quorum, pentapeptide

## Abstract

The layer of peptidoglycan surrounding bacteria provides structural integrity for the bacterial cell wall. Many organisms, including human cells and diverse bacteria, detect peptidoglycan fragments that are released as bacteria grow. Uropathogenic *Escherichia coli* (UPEC) strains are the leading cause of human urinary tract infections (UTIs) and many patients experience recurrent infection after successful antibiotic treatment. The source of recurrent infections may be persistent bacterial reservoirs *in vivo* that are in a quiescent state and, thus, are not susceptible to antibiotics. Here, we show that multiple UPEC strains require a quorum to proliferate *in vitro* with glucose as the sole carbon source; at low density, the bacteria remain viable but enter a quiescent, non-proliferative state. Of all clinical UPEC isolates tested to date, 35% (51/145) enter this quiescent state, including archetypal strains CFT073 (from classic endemic lineage ST73) and JJ1886 (from recently emerged, multidrug-resistant pandemic lineage ST131). We further show that quorum-dependent UPEC quiescence is prevented and reversed by small molecules, called proliferants, that stimulate growth, such as L-lysine, L-methionine, and peptidoglycan (PG) stem peptides, including an isolated PG pentapeptide from *Staphylococcus aureus.* Together, our results indicate that (i) uptake of L-lysine and (ii) PG peptide sensing by UPEC modulate the quorum-regulated decision to proliferate and further demonstrate that PG fragments are important for intra- and interspecies signaling in pathogenic *E. coli*.

## Introduction

At least 80% of urinary tract infections (UTIs) are caused by uropathogenic *Escherichia coli* (UPEC) strains, and 27% of patients with a UTI experience recurrence within 12 months after successful antibiotic treatment^1–3^. UPEC appear to enter a quiescent, non-growing state within urothelial transitional cells in the bladder wall that may enable *E. coli*, long after antibiotic treatment has been terminated, to resume division within bladder urine after apoptotic release from those cells^4,5^.

Recently, we reported the discovery of a new *E. coli* physiological growth state, common to many UPEC strains^6^. That is, the UPEC strain *E. coli* CFT073, a member of the major UPEC lineage sequence-type (ST) 73, and approximately 80% of ST73 strains tested grow well in liquid glucose minimal medium, but become quiescent, i.e., fail to grow, but remain viable, on glucose minimal agar plates at a cell density of less than 10^6^ CFU per plate, yet grow well on glucose minimal agar plates seeded with >10^6^ CFU. Moreover, 23% of randomly selected UPEC strains of diverse STs isolated from community-acquired UTIs also become quiescent on glucose minimal plates seeded with ≤10^6^ CFU^6^. Remarkably, these results indicate that growth of many UPEC strains on glucose minimal agar plates is quorum dependent^6^. Alternative sole carbon sources, including acetate, arabinose, fructose, fucose, galactose, gluconate, *N*-acetylglucosamine (NAG), maltose, and mannose also lead to cells entering quiescence at low plating density, whereas glycerol, ribose and xylose promote robust growth^6^. We also reported that mini-Tn5 transposon insertions in five central carbon metabolism genes (*gdhA, gnd, pykF, sdhA*, and *zwf*) prevent quiescence, suggesting that metabolic function must remain intact during quiescence even though the cells are nonproliferative. *In vitro* CFT073 quiescence is prevented when cells are placed near a single colony of actively growing CFT073 or *E. coli* MG1655, a K-12 laboratory strain, suggesting that the actively growing cells release one or more molecules that act as a proliferant. Additionally, we found that L-lysine is a proliferant for quiescent CFT073 cells, but only in combination with either L-methionine or L-tyrosine. Notably, the remaining 17 amino acids play no role in preventing quiescence^6^ and CFT073 is not auxotrophic for lysine, methionine or tyrosine. The quorum-dependency of CFT073 proliferation implies that the ability to sense a cell-to-cell signal(s), which may include L-lysine, L-methionine and/or L-tyrosine, underlies the decision by each individual cell to grow and divide or to enter the quiescent state, suggesting that the proliferative/quiescent transition may be controlled by a quorum sensing system^7,8^.

Here we show that CFT073 quiescent cells are filamentous, that L-lysine must be transported into CFT073 to prevent quiescence, suggesting that L-lysine synthesis is inhibited in quiescent cells, and that peptidoglycan fragments, which are normally secreted by growing *E. coli*^9^, and, more specifically, a peptidoglycan stem pentapeptide, prevent quiescence of two UPEC strains, CFT073, a prototypic ST73 UPEC strain and JJ1886, a pandemic isolate that displays increased sensitivity to proliferants. While peptidoglycan fragments have previously been shown to regulate development of *Bacillus subtilis* and *Mycobacterium tuberculosis*, they have not been shown to act as quorum-signaling molecules in UPEC^10–12^. These findings demonstrate that population-sensing and metabolite availability modulate the quiescent state of UPEC. In addition, these studies further define an *in vitro* model system for UPEC quiescence that will enable closer examination of quiescent UPEC physiology and may lead to new approaches to prevent recurrent UTI.

## RESULTS

### Quorum-dependent growth of UPEC and prevention of quiescence on glucose minimal agar plates by secreted signals

UPEC strain CFT073 enters a quiescent state when plated at low cell density (10^4^ cells per plate) on glucose minimal agar, yet grows robustly when plated at high cell density (10^8^ cells per plate), demonstrating that growth of CFT073 on glucose minimal agar plates is quorum-dependent (Fig. S1A)^6^. As reported previously, quiescent CFT073 cells can be resuscitated by challenge with specific external stimuli, called proliferants, including additions of 5 μl of a solution containing L-lysine (1 mM) and L-methionine (1 mM), 5 μl of human urine, or a single colony of actively growing *E. coli* spp. (CFT073 or MG1655), added to the plate^6^. In a soft agar overlay containing 10^4^ quiescent CFT073 cells, we tested known proliferants and observed the development of a zone of actively growing colonies surrounding the site of challenge for all proliferants tested over 24 h (Fig. 1A). Without challenge, cells failed to grow throughout the course of the experiment and remained quiescent. We also tested additional clinical isolates and found that quorum-dependent growth on glucose minimal agar is common among the ST131 UPEC strains in this study (8/33) (Table 1) and among other isolates in a previous study ^6^; however, we discovered that the number of cells constituting a quorum varies among strains. For example, the ST131 strain JJ1886 was proliferative on glucose minimal agar at a plating density greater than 10^3^ CFU, but quiescent at or below 10^3^ CFU (Table 1). Of the 35 strains tested in this study, 10 were reversibly quiescent on glucose minimal agar, and three of those strains were quiescent at or below a plating density of 10^3^ CFU, but non-quiescent at 10^5^ CFU (Table 1). These results show that sensitivity for quorum detection varies by strain, i.e., is strain-specific.

**Fig. 1.**
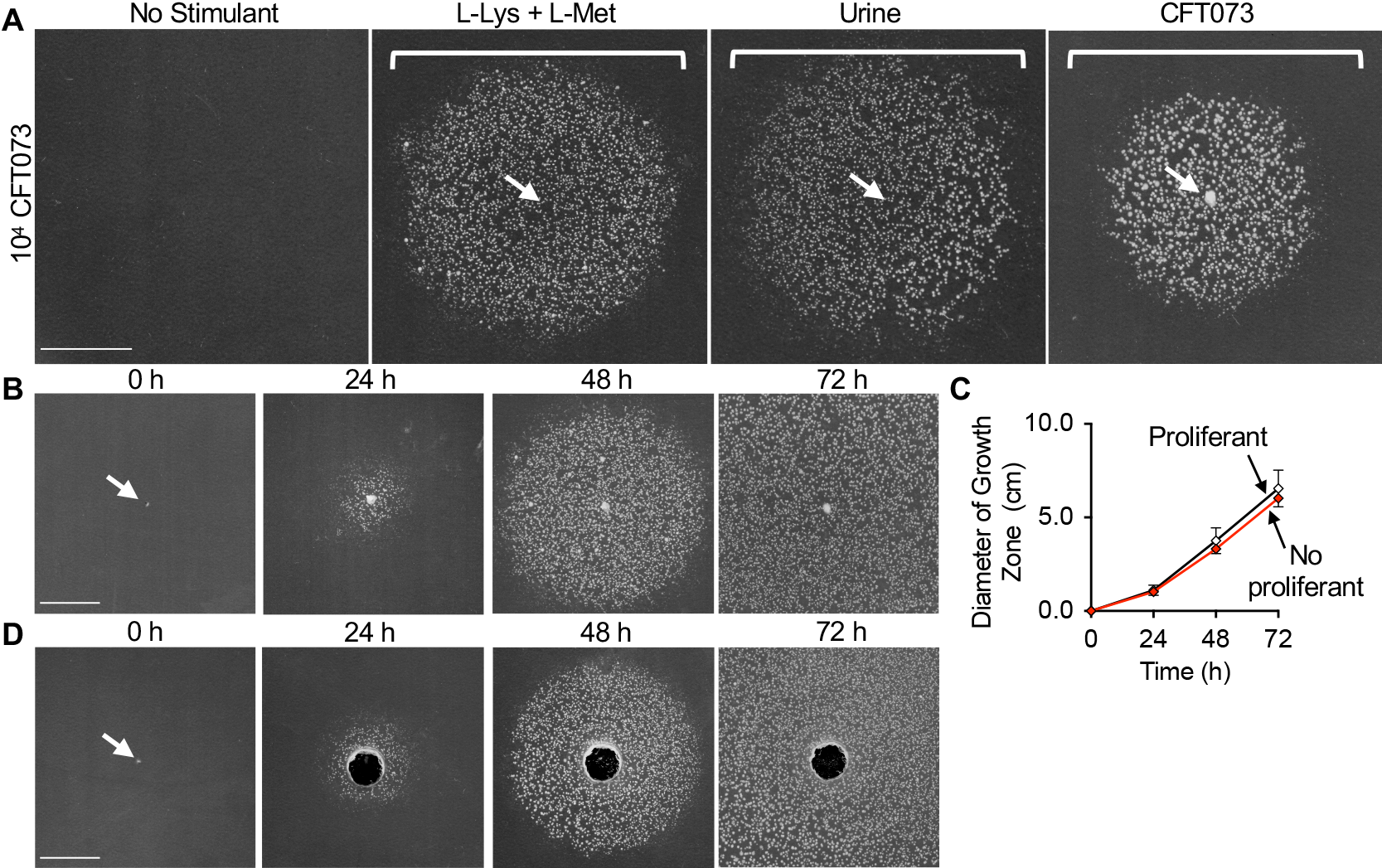
The quorum-regulated quiescent state of *E. coli* CFT073 cells on glucose M9 minimal agar is reversible. (A) CFT073 cells (10^4^) were plated onto M9 minimal agar with 0.2% glucose. A 5 μl solution of L-lysine (1 mM) and L-methionine (1 mM), sterile human urine, or an actively growing colony of CFT073 was transferred to the center of the plate, cells were incubated for 24 h, and the zone of growth surrounding the challenge site was monitored. (B, C) Zones of growth expand with time after addition of an actively growing colony of CFT073 up to 72 h. (D) Removal of the actively growing colony at 24 h has no effect on zonal expansion. White arrows indicate position of stimulant addition, brackets indicate the zone of colony growth and scale bars are 1 cm in length.

**TABLE 1.** Prevalence of quiescence among UPEC ST131 isolates

| Strain | Sequence type | Clade | Quiescent at 10<sup>3</sup> CFU<br>and reversed by Lys,<br>Met and Tyr <sup>a</sup> | Source, reference |
| --- | --- | --- | --- | --- |
| CFT073 | ST73 |  | + | 6 |
| Nissle 1917 | ST73 |  | + | 6 |
| JJ1886 | ST131 | C2 | + <sup>b</sup> | J. Johnson |
| JJ1887 | ST131 | C2 | - | J. Johnson |
| JJ2547 | ST131 | C2 | + <sup>b</sup> | J. Johnson |
| JJ2050 | ST131 |  | - | J. Johnson |
| JJ2528 | ST131 | C1 | - | J. Johnson |
| MVAST0036 | ST131 | C1 | - | J. Johnson |
| MVAST014 | ST131 | C1 | + <sup>b</sup> | J. Johnson |
| H17 | ST131 | B0 | - | J. Johnson |
| JJ1901 | ST131 |  | - | J. Johnson |
| JJ2087 | ST131 | B | - | J. Johnson |
| JJ1999 | ST131 | B | - | J. Johnson |
| MVAST167 | ST131 | A | - | J. Johnson |
| MVAST020 | ST131 | A | - | J. Johnson |
| JJ2134 | ST131 | C2 | + | J. Johnson |
| JJ2183 | ST131 | C2 | - | J. Johnson |
| JJ2489 | ST131 | C2 | - | J. Johnson |
| QUC02 | ST131 |  | - | J. Johnson |
| JJ2118 | ST131 | C1 | - | J. Johnson |
| JJ2193 | ST131 | C1 | - | J. Johnson |
| JJ2210 | ST131 | C1 | - | J. Johnson |
| JJ2578 | ST131 | C1 | - | J. Johnson |
| JJ2608 | ST131 | C1 | - | J. Johnson |
| MVAST038 | ST131 | C1 | - | J. Johnson |
| MVAST046 | ST131 | C1 | + | J. Johnson |
| MVAST077 | ST131 | C1 | - | J. Johnson |
| MVAST084 | ST131 | C1 | - | J. Johnson |
| MVAST131 | ST131 | C1 | - | J. Johnson |
| MVAST158 | ST131 | C1 | - | J. Johnson |
| MVAST179 | ST131 | C1 | + | J. Johnson |
| JJ2546 | ST131 |  | - | E. Sokurenko |
| JJ2548 | ST131 |  | + | E. Sokurenko |
| JJ2963 | ST131 |  | - | E. Sokurenko |
| JJ2974 | ST131 |  | + | E. Sokurenko |

<sup>a</sup> Represents strains seeded $\leq 10^3$ CFU in 0.2% glucose M9 minimal media 37 °C 24 h, (+) represents quiescent, (-) represents non-quiescent.
<sup>b</sup> Denotes strains that are non-quiescent when seeded at 10<sup>5</sup> CFU on 0.2% glucose M9 minimal media and grown at 37 °C for 24 h.

Next, we monitored the zone of growth of CFT073 cells as they exited quiescence over 72 h. First, we poured soft agar plates containing quiescent CFT073 cells seeded at 10^4^ CFU per plate, placed an actively growing colony of CFT073 onto the surface of each plate by toothpick and measured the zones of growth around the sites of inoculation at 24, 48 and 72 h. We observed that the zone of growth expands in diameter until the entire plate is covered by a lawn of actively growing cells (Fig. 1B and 1C). There are two potential scenarios to explain this result. First, the proliferant(s) that stimulates growth of quiescent cells is continuously released from the toothpicked colony throughout the experiment and induces growth of distal quiescent cells as it diffuses across the plate. Alternatively, the proliferant(s) that stimulates growth of quiescent cells is released from the toothpicked colony but, upon release of the signal, quiescent cells, once stimulated to proliferate, secrete their own signal, thereby stimulating adjacent quiescent cells to grow. To distinguish between these possibilities, glucose minimal agar plates were seeded with 10^4^ quiescent CFT073 cells from a stationary phase liquid glucose minimal medium culture, and a single CFT073 colony was toothpicked onto the center of each plate. The plates were then incubated for 24 h. During that time, a small zone of growth formed on each plate around the central toothpicked colony (Fig. 1B, 1C and 1D). After the zone had formed at 24 h, the toothpicked colony was removed to stop the colony from secreting additional signal(s). Even after removal of the toothpicked colony from the plate, the zone of growth continued to expand, unabated, for an additional 48 h and at a similar rate as the control plate, where the colony had been left intact (Fig. 1C and 1D). These results suggest that the proliferation-promoting signal propagates across the plate as CFT073 cells exit quiescence. These results further show that quiescent CFT073 cells at the edge of the plate remain viable and can be stimulated to proliferate for up to 72 h.

### Quiescent CFT073 cells are filamentous on glucose minimal agar plates

During uropathogenic infection, bacterial cells from quiescent intracellular reservoirs have several different morphologies, including rods, spheres, and filaments^13–15^. To determine if quiescent cells in our *in vitro* quiescence system also display morphological features that are distinct from actively growing cells, we compared the cell morphologies of CFT073 cells under several culture conditions, including LB agar and glucose minimal agar supplemented with L-lysine (1 mM) and L-methionine (1 mM), where indicated (Fig. 2A). After 24 h on LB agar or on glucose minimal agar supplemented with L-lysine and L-methionine, colonies were apparent and individual cells collected appeared as short rods with average lengths of 1.72 ± 0.59 μm (n = 200) and 2.33 ± 0.80 μm (n = 200), respectively (Fig. 2A and 2B). However, after 24 h on glucose minimal agar plates, no colonies were detected and quiescent cells harvested from the plate had a broad length distribution with a mean length of 7.17 ± 4.34 μm (n = 200), which is 300% longer than cells grown on glucose minimal agar supplemented with lysine and methionine (Fig. 2A and 2B). Many quiescent cells were longer than 10 μm and 16.5% of the cells appeared to contain one or more incomplete septa (Fig. 2A and 2B) (Fig. S1B). To determine if cell length changes upon extended incubation, we also measured cell length at 0 h and 48 h after entry into quiescence. At 0 h, cells harvested from a stationary phase culture grown in liquid glucose minimal medium were short rods with an average cell length of 2.14 ± 0.58 μm (n = 200). After 48 h on glucose minimal agar plates, quiescent cells were filamentous with an average cell length of 6.84 ± 3.76 μm (n = 72), which is similar to the average length at 24 h (Fig. S1B and S1C). These results suggest that quiescent CFT073 cells elongate during the first 24 h on the plate but fail to complete division.

**Fig. 2.**
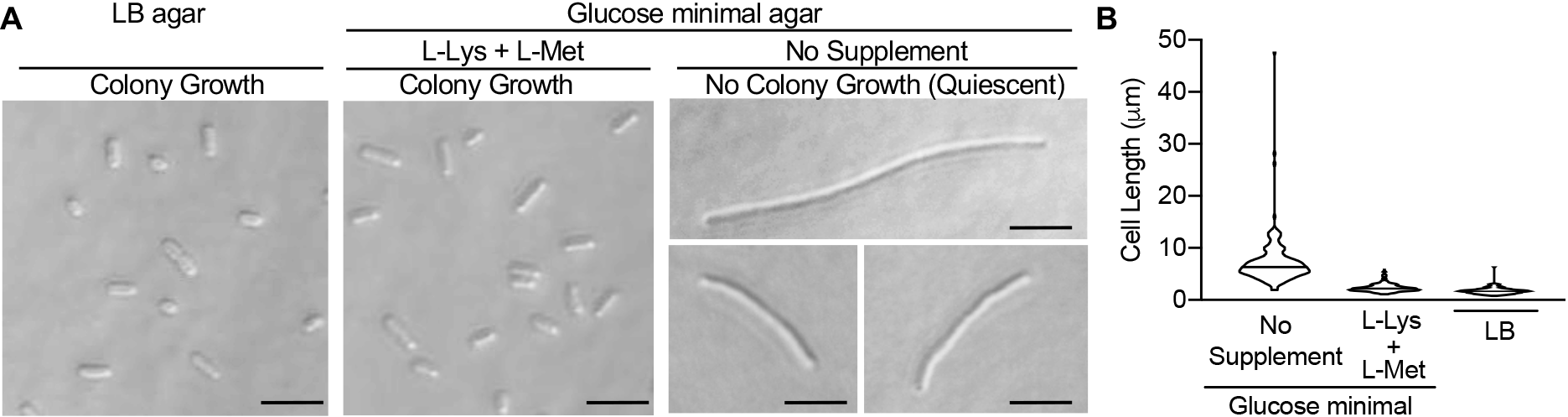
Quiescent CFT073 cells are filamentous at 24 h. (A) CFT073 cells were grown on LB agar or 0.2% glucose minimal agar with and without L-lysine (1 mM) and L-methionine (1 mM), harvested at 24 h and visualized by DIC microscopy. Scale bars are 5 μm in length. (B) Violin plot representing cell length distribution at 24 h for 200 cells grown under conditions described in (A).

### Lysine import is necessary for lysine to prevent quiescence, but an additional signal that induces proliferation is released from actively growing cells

L-lysine is a proliferant for quiescent CFT073 cells on glucose minimal agar containing L-methionine. To determine whether L-lysine must be imported into CFT073 cells to prevent quiescence, we constructed a CFT073 *lysP* insertion-deletion mutant by λ-Red recombination (Table S1). Of the three known lysine importers in *E. coli*, LysP is the major importer and a member of the amino acid, polyamine and organocation (APC) family of transporters^16,17^. CFT073 Δ*lysP* cells grew well overnight in liquid glucose minimal medium, similar to the wild type CFT073 strain (Fig. S2A) and exhibited quiescence on glucose minimal agar at low cell density (10^4^ cells per plate) (Fig. 3); yet, it grew robustly when plated at high density (10^8^ cells per plate). Unlike wildtype CFT073, when 5 μl of a solution of L-lysine (1 mM) and L-methionine (1 mM) was added to quiescent CFT073 *DlysP* cells plated at 10^4^ CFU, no colony growth was observed after 24 h of incubation suggesting that LysP-dependent transport is critical for lysine to stimulate proliferation. This suggests that lysine biosynthesis is inhibited in quiescent cells. CFT073 *DlysP* was complemented with a plasmid containing *lysP* (pLysP), which restored the ability of the strain to proliferate upon addition of L-lysine in the presence of L-methionine at low cell density (Fig. 3). Although quiescent CFT073 Δ*lysP* cells failed to respond to L-lysine, they proliferated after a colony of actively growing CFT073 cells was added to the plate (Fig. 3). In addition, CFT073 Δ*lysP* quiescent cells also proliferated when 5 μl of human urine was added to the plate (Fig. 3). These results suggest that import of L-lysine is required for L-lysine to stimulate proliferation and override the absence of the quorum. Moreover, one or more L-lysine-independent proliferants that reverse quiescence are secreted from bacterial colonies and are present in urine.

**Fig. 3.**
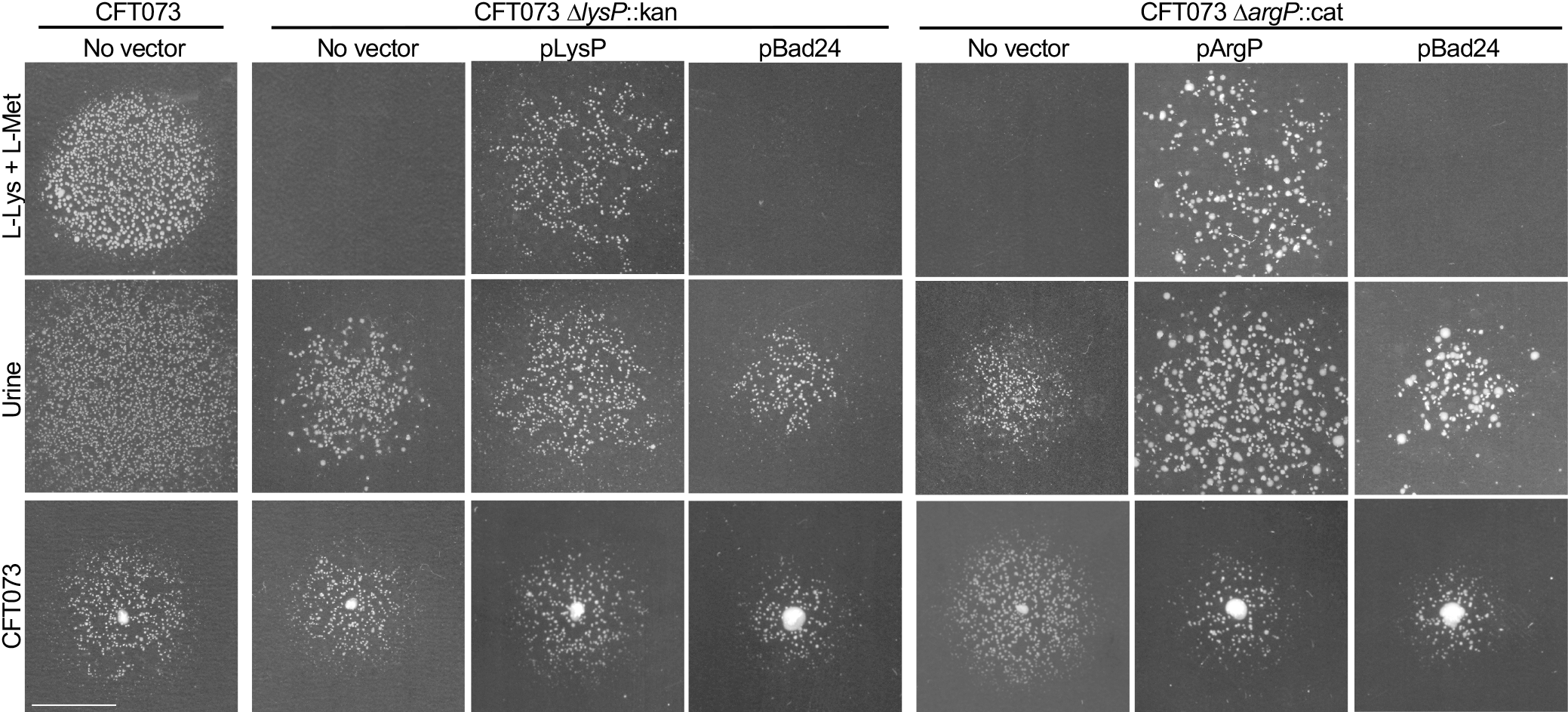
Quiescent CFT073 Δ*lysP* and Δ*argP* cells fail to proliferate with L-lysine. CFT073, CFT073 Δ*lysP*, CFT073 Δ*lysP* with pLysP, CFT073 Δ*lysP* with pBad24, CFT073 Δ*argP*, CFT073 ΔargP with pArgP and CFT073 ΔargP with pBad24 cells (10^4^) were plated onto M9 minimal agar with 0.2% glucose. All vector carrying strains were plated onto agar containing 50 μg/ml Ampicillin, while CFT073 Δ*lysP* with pLysP and CFT073 Δ*lysP* with pBad24 were plated onto agar with 0.1% arabinose. Quiescent cells were challenged with a 5 μl solution of L-lysine (0.5 mM) and L-methionine (1 mM) (CFT073 Δ*lysP*, CFT073 Δ*lysP* with pLysP, CFT073 Δ*lysP* with pBad24), L-lysine (1 mM) and L-methionine (1 mM) (CFT *ΔargP*, CFT073 Δ*argP* with pArgP and CFT073 ΔargP with pBad24), sterile human urine or an actively growing colony of MG1655 transferred to the center of the plate and cells were incubated for 24 h.

ArgP is a transcriptional regulator of *lysP*^16,18^. In the absence of lysine, ArgP positively regulates transcription of *lysP*, as well as several *E. coli* genes involved in lysine and peptidoglycan biosynthesis, including *dapB* (dihydropicolinate reductase), *dapD* (tetrahydrodipicolinate succinylase), *lysA* (diaminopimelate decarboxylase) and *lysC* (aspartokinase III)^16,18–20^. To test whether ArgP plays a regulatory role in quorum-dependent growth or quiescence, we deleted *argP*. Like wild type CFT073, CFT073 Δ*argP* cells grew well overnight in liquid glucose minimal medium (Fig. S2A) and were quiescent at low cell density (10^4^ cells per plate) on glucose minimal agar (Fig. 3) indicating that ArgP is not essential for inducing quiescence at low density (Fig. 3). Similar to the result with CFT073 Δ*lysP* cells, quiescent CFT073 Δ*argP* cells failed to respond to the addition of L-lysine with L-methionine after 24 h of incubation, but the cells proliferated in response to a colony of CFT073 toothpicked to the center of a glucose minimal agar plate seeded with 10^4^ CFU CFT073 Δ*argP* (Fig. 3). Complementation of CFT073 *DargP* with the wildtype CFT073 *argP* gene on a plasmid (pArgP) restored the ability of the strain to respond to L-lysine in the presence of L-methionine at low cell density (Fig. 3). Since LysP expression has been reported to be repressed 35-fold in *E. coli argP* mutants^20^, this result supports the conclusion that external L-lysine must be transported into CFT073 via LysP in order to prevent quiescence and confirms that expression of LysP is regulated by ArgP. Moreover, these results also show that there is another molecule secreted by actively growing cells that stimulates proliferation of quiescent CFT073 cells.

To further understand why the population that defines a quorum varies between strains, we investigated the ST131 clinical isolate JJ1886, which exhibits quiescence on glucose minimal agar at or below a plating density of 10^3^ CFU but grows robustly at 10^5^ CFU and in liquid glucose minimal medium (Fig. 4A and S2B). We observed that 5 μl of a solution of L-lysine (1 mM) alone is sufficient to promote growth and prevent quiescence of JJ1886 plated at low density on glucose minimal agar (10^3^ CFU per plate), although growth is more efficient in the presence of L-methionine (1 mM), and L-methionine (1 mM) alone does not stimulate growth (Fig. 4B). Since quiescent JJ1886 cells on glucose minimal agar proliferate in the presence of L-lysine alone, we titrated the L-lysine concentration to determine the threshold for stimulation of growth and compared JJ1886 to CFT073. We determined that a low concentration of L-lysine (5 μl of 0.1 mM) is sufficient to promote proliferation of quiescent JJ1886 cells. In contrast, quiescent CFT073 cells only proliferate at a higher concentration of L-lysine (5 μl of 1 mM) and only in the presence of L-methionine. These results suggest that although JJ1886 behaves phenotypically similar to CFT073 with respect to quiescence, it is more sensitive to detecting or responding to an external signal(s) that prevents quiescence. We then tested if quiescent JJ1886 also responds to the other known CFT073 proliferants, including 5 μl of urine or a colony of actively growing *E. coli* cells, and we observed that both were effective for stimulating proliferation of quiescent JJ1886 cells (Fig. 4B).

**Fig. 4.**
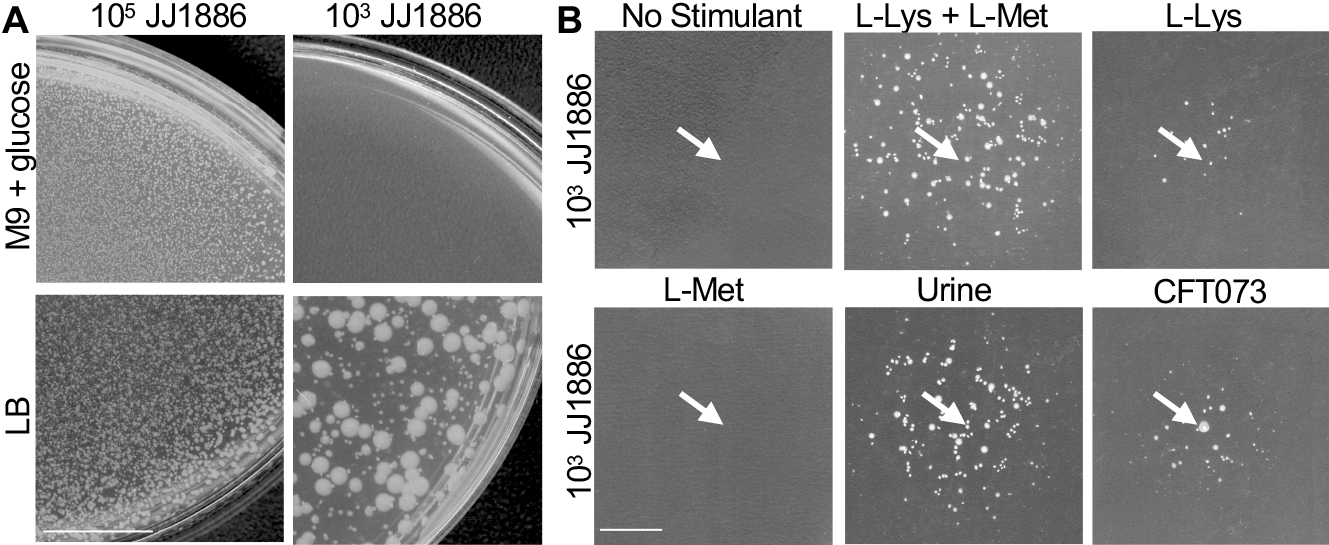
ST131 strain JJ1886 displays quorum dependent growth. (A) JJ1886 cells were plated onto M9 minimal agar with 0.2% glucose and LB agar, at 10^5^ and 10^3^ CFU. JJ1886 displays quorum dependent growth at 10^3^ CFU on glucose M9 minimal agar. (B) JJ1886 cells (10^3^) were plated onto M9 minimal agar with 0.2% glucose. A 5 μl solution of L-lysine (1 mM) and L-methionine (1 mM), L-lysine (1 mM), L-methionine (1 mM), sterile human urine, or an actively growing colony of CFT073 was transferred onto a JJ1886 seeded plate and cells were incubated for 24 h.

Since L-lysine stimulates proliferation of both JJ1886 and CFT073, we tested 5 μl of D-lysine (1 mM) and 5 μl of the biosynthetic lysine precursor diaminopimelic acid (Dap) (1 mM) for stimulating proliferation of quiescent cells. Both failed to stimulate growth of quiescent CFT073 and JJ1886 (Fig. S3A and S3B).

### Peptidoglycan fragments prevent UPEC quiescence

Next, we considered peptidoglycan (PG) as a potential signal that would communicate a quorum since peptidoglycan fragments, including di-, tri-, and tetrapeptides, are normally secreted by actively growing bacterial cells including *E. coli*^9,11,21,22^ We tested if peptidoglycan fragments from *E. coli* could prevent quiescence and stimulate proliferation of CFT073 and JJ1886. Peptidoglycan was digested with mutanolysin, a mixture of two N-acetylmuramidases and an N-acetylmuramyl-L-alanine amidase, to release a heterogeneous mixture of peptidoglycan fragments, including various muropeptides and stem peptides (Fig. 5A). The addition of the total mixture of digested peptidoglycan fragments from *E. coli* induced proliferation of quiescent JJ1886 cells, plated at 10^3^ CFU per plate (Fig. 5A), and quiescent CFT073 cells, plated at 10^4^ CFU per plate (Fig. S4A). Titration of the amount of total peptidoglycan added to the plate revealed dose-dependent detection of fragments leading to JJ1886 colony growth in a zone around the site of addition (Fig. S4B). Digested peptidoglycan from *Bacillus subtilis* and *Staphylococcus aureus* also stimulated proliferation of quiescent JJ1886 and CFT073 cells (Fig. S4C and S4D). Furthermore, peptidoglycan fragments were also effective for stimulating quiescent CFT073 Δ*lysP* cells to proliferate, indicating that the peptidoglycan response is independent of lysine import (Fig. S5A). Interestingly, these results show that quiescent *E. coli* cells respond to peptidoglycan fragments from various bacterial species, which can differ in the identity of the amino acid at the third position of the stem peptide, usually *meso-Dap* (i.e., *E. coli*) or L-lysine (i.e., *S. aureus*) ^11,23,24^.

**Fig. 5.**
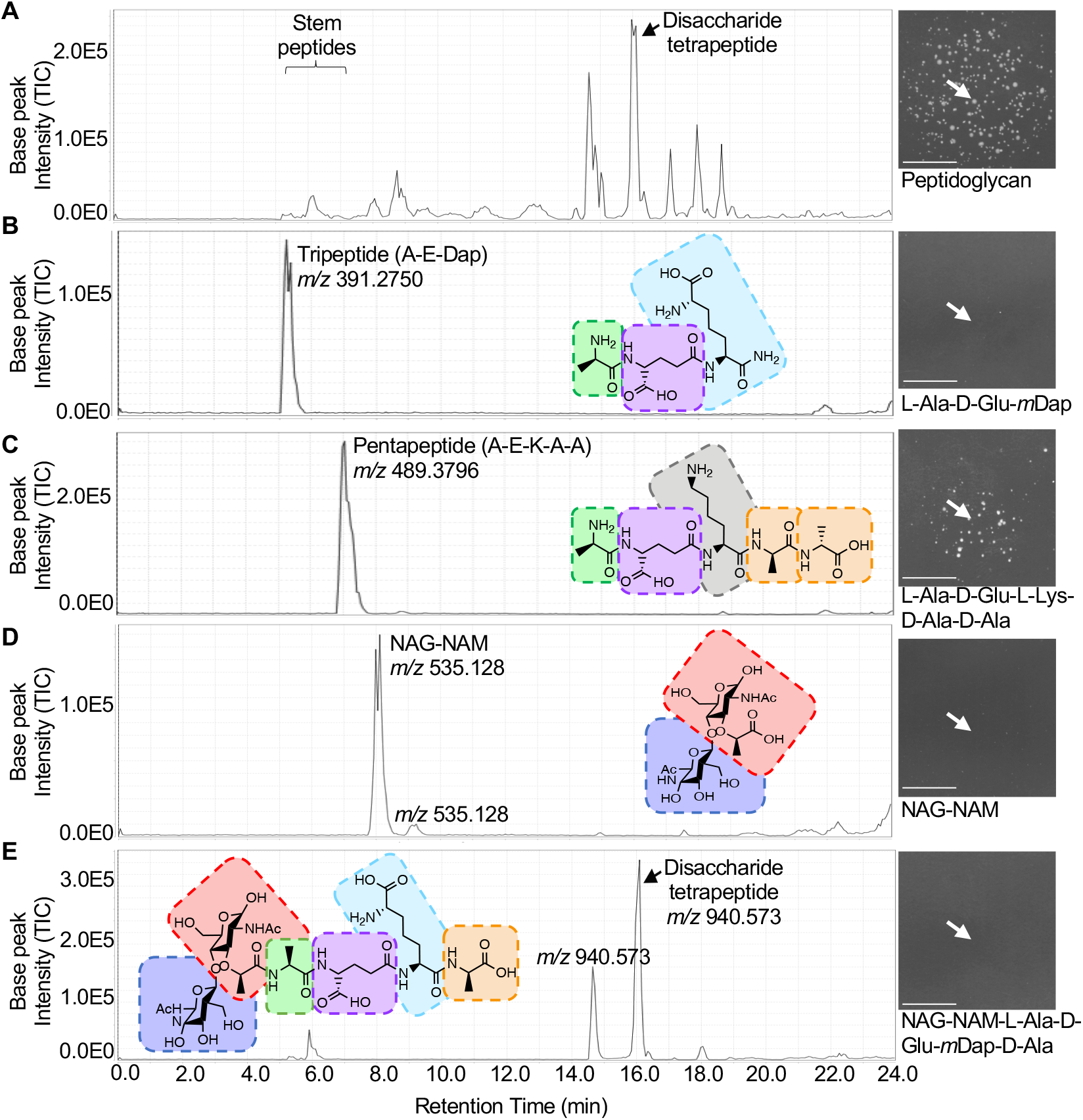
Peptidoglycan fragments promote proliferation of CFT073 and JJ1886 cells. LCMS profiles of total ion counts (TIC) including (A) mutanolysin digested crude PG, which was tested for stimulating proliferation of quiescent JJ1886 cells plated at 10^3^ CFU on M9 minimal agar with 0.2% glucose, inset; (B) elution of a PG tripeptide similar to an *E. coli* fragment and corresponding test for quiescence reversal, inset; (C) elution of a PG pentapeptide similar to a *S. aureus* fragment and corresponding test for quiescence reversal, inset; (D) elution of a NAG-NAM disaccharide and corresponding test for quiescence reversal, inset; and (E) elution of a disaccharide tetrapeptide isolated from the mutanolysin-digested crude PG corresponding test for quiescence reversal, inset. Candidate proliferants were collected tested in the quiescence assay as described in *Material and Methods* and plates were incubated at 37°C for 24 h. White arrows indicate position of proliferant addition and scale bar is 1 cm in length.

### A peptidoglycan stem pentapeptide acts as a signal to promote proliferation of quiescent JJ1886 and CFT073 cells

To identify which fragment(s) of digested peptidoglycan stimulates proliferation of quiescent cells, we performed high performance liquid chromatography-mass spectrometry (HPLC-MS) on *E. coli* peptidoglycan fragment mixtures (Fig. 5A) and tested fractions across the elution for the ability to promote proliferation of quiescent JJ1886 cells. We found that early fractions collected, with retention times up to 10 minutes were sufficient to stimulate proliferation of quiescent JJ1886 cells (Fig. S5B). Tandem mass spectrometry confirmed that these fractions contain low molecular weight peptidoglycan fragments, consistent with stem peptides, such as the tri-and tetrapeptides.

To further understand the molecular determinants of PG peptides that stimulate proliferation of quiescent cells, we compared several highly pure stem peptide fragments of varying amino acid compositions and lengths. We observed that a tripeptide, representing the minimal unit of an *E. coli* peptidoglycan stem peptide that could be generated from amidase and carboxypeptidase activities in the *E. coli* periplasm (L-Ala-D-Glu-*m*Dap), failed to stimulate proliferation of quiescent JJ1886 cells (Fig. 5B). However, a peptidoglycan pentapeptide, L-Ala-D-Glu-L-Lys-D-Ala-D-Ala (A-E-K-A-A), corresponding to the stem pentapeptide present in *S. aureus* peptidoglycan, stimulated proliferation of quiescent JJ1886 (Fig. 5C) and CFT073 (Fig. S5A) cells. We also tested a smaller fragment of the pentapeptide, acetyl-L-Lys-D-Ala-D-Ala (K-A-A) but did not observe stimulatory activity (Fig. S6A). Fragments including the disaccharide moiety alone, containing NAG and N-acetyl muramic acid (NAM) (NAG-NAM) (Fig. 5D), and higher molecular weight molecules with the attached stem peptide (i.e., disaccharide tetrapeptide) (Fig. 5E), in various states of crosslinking, eluted later, after 10 min, and had no proliferant activity (Fig. S7). NAG-NAG, NAG and NAM were tested, but none of these sugars stimulated proliferation of quiescent cells (Fig. S6B). These results further suggest that a stem peptide functions as a signal to promote proliferation of quiescent cells.

Finally, to further clarify if a stimulatory peptide, such as the pentapeptide, exerts its function without entering the cytoplasm or if it is imported into a quiescent cell and enters the peptidoglycan recycling pathway, we constructed JJ1886 strains deleted for the oligopeptide transporter operon, *oppABCDF*^25–29^, as well as the peptidoglycan tripeptide-specific delivery protein *mppA*^27,30–32^. Both deletion strains grew well in liquid glucose minimal medium and became quiescent when plated at 10^3^ CFU per plate, similar to the parental strain JJ1886; however, both strains were stimulated to proliferate upon addition of the pentapeptide to the plate (Fig. S8), indicating that it is not necessary for the pentapeptide to enter the cytoplasm via the Opp transporter to prevent quiescence and therefore it may act as a signal to prevent and reverse quiescence.

## DISCUSSION

Here, we show that two virulent UPEC strains, CFT073 and JJ1886, enter a quiescent state when plated on glucose minimal agar that is prevented and reversed by proliferants, including a pentapeptide derived from PG, and L-lysine. Quiescence is a unique physiological state recently described for *E. coli* that is common among UPEC strains (51/145 isolates tested) and regulated by the presence or absence of a quorum. Several notable features of the *E. coli* quiescent phenotype are described here. First, the quiescent state of UPEC appears to be a bona fide developmental state that is adopted by an entire population of cells under a given growth condition. This is in contrast to persistence, which occurs in a small percentage of cells^33–37^. Second, adoption of a non-proliferative state would require that many physiological systems would be slowed or inhibited, including DNA replication, transcription, translation, cell division and peptidoglycan synthesis. Accordingly, we observed that although quiescent cells adopt a filamentous morphology, they fail to divide or successfully proliferate upon extended incubation (Fig. S1B). Furthermore, as described previously, mutations in five central carbon metabolism genes (*gdhA, gnd, pykF, sdhA*, and *zwf*) prevent quiescence and promote growth, indicating that specific metabolic pathways must be functional to maintain the quiescent state^6^. Finally, different strains respond to known proliferants with varying sensitivities, suggesting that either sensitivity of detection or regulatory control over the physiological systems varies among strains.

Peptidoglycan muropeptide fragments are known to resuscitate dormant *Mycobacterium smegmatis*^38^ and to signal *Bacillus subtilis* endospores to germinate^12^. Here in *E. coli*, peptidoglycan fragments not only cause cells to reenter the cell cycle, but they may also act as a signal to communicate the presence of a quorum, although they do not appear to require uptake by the Opp transport system. Previously, it was shown that an *oppA* mutant is far less infective in a mouse UTI model than the wildtype^39^, and it was suggested that small oligopeptides, present in high concentration in urine^39^, are imported and serve as major nutrients for CFT073 growth in urine^39^. Here, the pentapeptide does not enter the quiescent cells via Opp. Therefore, either the pentapeptide is imported by a second, presently unknown, oligopeptide permease or the pentapeptide acts as a signal in the periplasm and is recognized by a membrane receptor or signal transduction system that, when activated, allows cell proliferation. The serine/threonine kinase PknB is a widely conserved peptidoglycan binding protein that is important for germination of spores in *B. subtilis*^12^; however, no similar proteins have been found thus far in *E. coli*.

The cause(s) of recurrent UTIs are complex^40^, however, it appears that UPEC can bind to, enter, and replicate within superficial facet cells in the mouse and human bladder epithelium, resulting in intracellular biofilm-like communities (IBCs)^40,41^. IBCs escape from infected superficial facet cells within hours of development and several cycles of IBC formation occur, but each successive round is associated with a reduced replication rate and smaller IBCs^15^. The infected superficial facet cells exfoliate, exposing underlying transitional epithelial cells that become infected with IBC-derived UPEC progeny. Upon infection of transitional cells, small numbers of UPEC appear to enter a latent or quiescent intracellular state in endosomal vesicles (2 to 12 cells/vesicle), establishing what have been termed quiescent intracellular reservoirs (QIRs)^42^ resistant to antibiotic treatment. Our results from *in vitro* quiescence assays with UPEC described here suggest that upon release from endosomal vesicles, UPEC may sense proliferants and thus be stimulated to reenter the cell cycle.

## MATERIALS AND METHODS

### Bacterial strains, plasmids and growth conditions

*E. coli* strains and plasmids used in this study are listed in Table 1. Clinical UPEC isolates were obtained from the collections of J. Johnson (29 strains) and E. Sokurenko (5 strains) (Table 2). LB agar (Lennox) was used for routine cultivation. Amino acids and M9 minimal medium and agar with glucose (0.2% or 0.4%, where indicated) were prepared as described by^6^. Gene deletions were constructed by λ-Red recombination as described in Datsenko and Wanner and confirmed by sequencing^43^. Culture density was monitored by growing cells in liquid 0.2% glucose M9 minimal medium and measuring OD_600_ at the indicated times.

### Complementation

Plasmids for complementation of both CFT073 *ΔlysP* and CFT073 *ΔargP* gene deletions were constructed using expression vector pBAD24. Both genes were amplified from CFT073 by whole colony PCR. The resulting PCR products were digested with EcoRI and HindIII. Digested PCR products were ligated into EcoRI/HindIII-digested pBad24. Both pLysP and pArgP plasmid constructs were verified by sequencing and then transformed into either CFT073 *ΔlysP* or CFT073 *ΔargP* deletion strains by electroporation. To test for complementation, transformants were grown overnight in 0.2% glucose M9 minimal medium with ampicillin (50 μg ml^−1^) and assayed for quiescence on glucose minimal agar supplemented with arabinose (0.1%), where indicated.

### Quiescence assay

Quiescence assay was performed as described previously^6^. CFT073 and JJ1886 wild type and mutant strains were grown on LB agar plates overnight at 37 °C. An isolated colony was used to inoculate 5 ml of 0.2% glucose M9 minimal media, incubated at 37 °C with shaking overnight, and then diluted to between 10^3^ to 10^8^ CFU, as indicated, in 3 ml 0.2% glucose M9 minimal medium containing 0.75% Difco noble agar held at 45 °C. Inoculated soft agar was immediately poured onto plates containing a 12 ml layer of 0.2% glucose M9 minimal medium with 1.5% noble agar or directly into an 8.5 cm^2^ polystyrene culture dish. Proliferants, including actively growing cells, amino acids, Dap (2,6-diaminopimelic acid), sterile human urine and peptidoglycan, were added as indicated and plates were incubated at 37 °C between 24 and 72 h and then imaged.

### Microscopy

Overnight cultures of CFT073 were grown to stationary phase in 0.2% glucose M9 minimal media. Five μl of a 3-log diluted culture was spotted onto LB agar or 0.2% glucose M9 minimal medium with 1.5% noble agar and, where indicated, supplemented with L-lysine (1 mM) and L-methionine (1 mM), then allowed to dry for 1 h at room temperature. Plates were incubated at 37 °C for 24 or 48 h. Cells were harvested onto a glass cover slip, resuspended in phosphate-buffered saline (PBS) with ethylenediaminetetraacetic acid (EDTA) (1 mM) and applied to an agar gel pad containing 0.2% glucose M9 minimal with 0.5% noble agar. Cells were visualized by differential interference contrast (DIC) microscopy using a Zeiss LSM 700, and images were captured on an AxioCam HRc high-resolution camera with ZEN 2012 software. Images were processed using Adobe Photoshop CC and analyzed using NIH ImageJ software.

### Peptidoglycan fragment separation

*E. coli* CFT073 was inoculated in 10 ml of M9 minimal medium containing 0.4% glucose and incubated at 37 °C shaking (200 rpm) for 24 h. Cultures were subsequently inoculated into one liter of M9 minimal media containing 0.4% glucose and incubated for 24 h shaking (175 rpm). Methods used for peptidoglycan isolation were adapted from Desmarais *et al.*, 2014^44^. Cells were harvested at 5,000 × *g* for 10 minutes at room temperature and pellets were resuspended in 3 ml of M9 minimal medium. Cell suspensions were transferred into 6 ml of 6% sodium dodecyl sulfate (SDS) and boiled for 3 h with stirring. After 3 h, heat was discontinued, and cell suspensions were stirred overnight. Cell suspensions were centrifuged at 262,000 × *g* for 30 minutes, pellets were resuspended in liquid chromatography mass spectrometry (LCMS) grade water and further centrifugation and wash steps occurred until no SDS remained. Final pellet was resuspended in 900 μl of 10 mM Tris-HCl with 0.06% w/v NaCl and 100 μg of Pronase E was added to the cell suspension and then incubated for 2 h at 60 °C. After incubation, 200 μl of 6% SDS was added and the mixture was boiled at 100 °C for 30 min. Samples were centrifuged at 262,000 × *g* for 30 min washed 4 times with LCMS grade water. The final pellet was resuspended in 200 μl of PBS and incubated with 200 U of mutanolysin (Sigma-Aldrich) at 37 °C overnight. Samples were centrifuged at 16,000 × *g* for 10 min to remove cellular debris, peptidoglycan supernatant was collected for further testing. Mutanolysin was removed by ultrafiltration using a polyethersulfone filter with a 3 kDa molecular weight cut-off.

Analytical high-performance liquid chromatography (HPLC) was used to purify the peptidoglycan extract. HPLC experiments were performed on a Shimadzu prominence-I LC-2030C equipped with a PDA detector model LC-2030/2040, pump LC-2030, and autosampler LC-2030. Preparative HPLC experiments were completed with a Waters XBridge Analytical C18 5 μm, 4.6 × 250 mm column at 30 °C under a solvent gradient of 10-100% B (5mM ammonium acetate with 15% acetonitrile pH 4.89) run over 16 minutes against solvent A (5mM ammonium acetate pH 4.37). Solvent B was held at 10% for 4 minutes prior to gradient beginning and 100% B was held for a subsequent 4 minutes. Individual peaks were collected according to 205 nm absorbance values. Initial chemical characterization was carried out by liquid chromatography mass spectrometry (LCMS) on a Thermo Fisher ISQ EC coupled to a Shimadzu Prominence-i LC-2030 system. Chemical separation utilized the same chemical gradient as previously with a reduced flow rate of 0.6 ml min^−1^. Chemical characterization of pure compounds was undertaken using electrospray ionization mass spectrometry (ESIMS) using an AB Sciex TripleTOF 4600 spectrometer and LC-MSMS experiments.

LC-MSMS methods were performed on a AB Sciex TripleTOF 5600 spectrometer coupled to a Waters H class UPLC. LC methods mirrored that of initial screening with an adjusted flow rate of 0.5 mL/min on an XBridge Analytical C18 5 μm, 4.6 × 250 mm column at room temperature. SWATH acquisition was performed in positive ionization mode. The method specific parameters were as follows: gas 1 (GS1) 55 psi, gas 2 (GS2) 60 psi, curtain gas (CUR) 25 psi. The source-specific parameters were: temperature (TEM) 500°C, ion spray voltage floating (ISVF) 5500 V, declustering potential (DP): 100, collision energy (CE) 10, collision energy spread (CES) 15. An initial TOF scan was collected from *m/z* 100 to 1500 and SWATH data was acquired (*m/z* 200-1100) over 36 SWATH windows per cycles with a window size of *m/z* 25 and a *m/z* 1 overlap between windows.

Peptidoglycan-derived stem peptide fragments including ‘A-E-K-A-A’ pentapeptide (L-Ala-D-Glu-L-Lys-D-Ala-D-Ala (Sigma-Aldrich), ‘K-A-A’ tripeptide (acetyl-L-Lys-D-Ala-D-Ala) (Sigma-Aldrich) and ‘A-E-Dap’ tripeptide (L-Ala-D-Glu-mDap) (AnaSpec), were each tested at 5 μl (2 mM) against strains JJ1886, JJ1886 Δ*oppABCDF* and JJ1886 Δ*mppA* (10^3^) grown on M9 minimal agar with 0.2% glucose for 24 h at 37 °C.

## Supplemental information

**Table S1**

**Figure S1, S2, S3, S4, S5, S6, S7, S8**

## ACKNOWLEDGEMENTS

We thank Cathy Trebino, Evelyn Siler, Josiah Morrison, Negar Rahmani, and Colby Ferreira for critical reading of the manuscript, Rebecca Dickinson for technical assistance, Barry Wanner for pKD267, Tyrrell Conway for helpful discussions, Rod Welch for CFT073 comparison strain, Evgeni Sokurenko for UPEC strains and plasmids, and Janet Atoyan for microscopy and sequencing assistance. Microscopy and sequencing were performed at the Rhode Island Genomics and Sequencing Center, which is supported in part by the National Science Foundation (MRI Grant No. DBI-0215393 and EPSCoR Grant No. 0554548 & EPS-1004057), the US Department of Agriculture (Grant Nos. 2002-34438-12688, 2003-34438-13111, and 2008-34438-19246) and the University of Rhode Island.

