## Supplemental information for "Peptidoglycan sensing prevents quiescence and promotes quorum-independent growth of uropathogenic *Escherichia coli*"

**TABLE S1. *E. coli* strains and plasmids used for genetic analyses and constructions in this study**

| *E. coli* Strain or Plasmid | Genotype | Identifier | Source, reference or construction |
|---|---|---|---|
| CFT073 Str$^r$ | Spontaneous streptomycin-resistant mutant of CFT073 | CFT073 | 6 |
| MG1655 | *LAM-rph-1* | MG1655 | 45 |
| BL21 (λde3) | F- ompT gal dcm lon hsdSB(rB-mB-) λ(de3[lacI lacUV5-T7 gene 1 ind1 sam7 nin5]) | | EMD Millipore, USA |
| ED0052 | CFT073 Str$^r$ *ΔlysP*::*parE-kan* | CFT073 *ΔlysP* | CFT073; λred |
| ED0070 | CFT073 Str$^r$ *ΔargP*::*cat* | CFT073 *ΔargP* | CFT073; λred |
| JJ1886 | | JJ1886 | 46 |
| ED0118 | JJ1886 *ΔmppA*::*cat* | JJ1886 *ΔmppA* | JJ1886; λred |
| ED0131 | JJ1886 *ΔoppABCDF*::*cat* | JJ1886 *ΔoppABCDF* | JJ1886; λred |
| Plasmids | | | |
| pKD46 | *amp* | | 43 |
| pKD3 | *cat* | | 43 |
| pKD267 | *kan* | | B. Wanner[a] |
| pKD119 | *tet* | | 43 |
| pBAD24 | *amp* (expression vector) | | 47 |
| pLysP | *amp* $P_{ara}$::*lysP* | pLysP | This study |
| pArgP | *amp* $P_{ara}$::arg*P* | pArgP | This study |

[a] J. Teramoto, K. A. Datsenko, and B. L. Wanner, unpublished

### Supplemental Figure Legends

**Fig. S1. Quiescent CFT073 cells are filamentous at 48 h**

(A) CFT073 cells were cultured on 0.2% glucose M9 minimal agar at low density ($10^4$), which promotes quiescence, and at high density ($10^8$), which prevent quiescence and promotes proliferation, leading to a lawn of colony growth on the agar plate. (B) CFT073 cells were grown on 0.2% glucose M9 minimal agar at 37 °C, harvested at 0 and 48 h and visualized by DIC microscopy. Scale bars are 5 μm in length. Arrows indicate position of failed septa. (C) Violin plot representing cell length distribution at 0 h (n = 200), and 48 h (n = 72) grown under conditions described in (B).

**Fig. S2. UPEC strain growth in glucose M9 minimal media**

(A) CFT073, CFT073 *ΔlysP*, CFT073 Δ*argP* and (B) JJ1886 were grown in 0.4% glucose M9 media at 37 °C for 24 h with $OD_{600}$ measured every 2 h from 0 to 12 h and 22 to 24 h. All strains grow robustly overnight in glucose M9 minimal media.

**Fig. S3. D-lysine and diaminopimelic acid (Dap) fail to stimulate proliferation of CFT073 and JJ1886 cells on glucose M9 minimal agar**

(A) CFT073 ($10^4$) and (B) JJ1886 ($10^3$) cells were grown on 0.2% glucose M9 minimal agar and challenged with a solution of either (A) L-lysine (1mM) and L-methionine (1 mM), D-lysine (1 mM) and L-methionine (1 mM), Dap (1mM) and L-methionine (1 mM), (B) L-lysine (1mM), D-lysine (1 mM) or Dap (1 mM). Plates were incubated for 24 h and pictures were taken. Growth was observed on plates challenged with L-lysine and L-methionine or L-lysine. Arrows indicate position of stimulant addition. Scale bars are 1 cm in length.

**Fig. S4. *E. coli*, *B. subtilis* and *S. aureus* peptidoglycan fragments stimulate proliferation of CFT073 and JJ1886 cells**

(A, C) CFT073 ($10^4$) and (D) JJ1886 ($10^3$) cells were plated onto M9 minimal agar with 0.2% glucose and challenged with a mixture of digested (A) CFT073 peptidoglycan (C, D) *B. subtilis* and *S. aureus* peptidoglycan or (A) a solution of PBS. Plates were incubated (D) 24 h and (A, C) 48 h. (B) Digested *E. coli* peptidoglycan was titrated (0, 5 and 10 µg challenged on JJ1886 seeded plates and CFU per zone was calculated. White arrows indicate position of stimulant addition, and scale bars are 1 cm in length.

**Fig. S5. Peptidoglycan fragments, including a *S. aureus* pentapeptide, stimulate proliferation of quiescent UPEC cells.**

(A) CFT073 ($10^4$), CFT073 *ΔlysP* and (B) JJ1886 ($10^3$) cells were plated onto M9 minimal agar with 0.2% glucose and challenged with a mixture of digested *B. subtilis* peptidoglycan (50 µg) or *S. aureus* pentapeptide (A-E-K-A-A) (5 µg). Plates were incubated for 48 to 72 h in (A). (B) Quiescent JJ1886 cells were challenged with early (4 to 10 min) or late (10 to 20 min) fractions of mutanolysin digested *E. coli* peptidoglycan fractionated by HPLC. Plates were incubated for 24 h. Arrows indicate position of proliferant addition and scale bars are 1 cm in length.

**Fig. S6. Candidate proliferants tested for reversal of JJ1886 quiescence**

LCMS profiles of total ion counts (TIC) for a PG tripeptide (A) similar to an *S. aureus* fragment, which was tested for stimulating proliferation of quiescent JJ1886 cells plated at $10^3$ CFU per plate on M9 minimal agar with 0.2% glucose, inset; (B) elution of a PG pentapeptide similar to a *S. aureus* fragment and corresponding test for quiescence reversal, inset; (C) Candidate proliferants including NAG (0.4 mM), NAM (0.4 mM) and NAG-NAG (1 mM) were tested for their ability to stimulate quiescent JJ1886 cells ($10^3$ per plate) on 0.2% glucose M9 minimal agar and plates were incubated for 24 h. White arrows indicate position of proliferant addition and scale bars are 1 cm in length.

**Fig S7. *E. coli* peptidoglycan fragment analysis by tandem mass spectrometry**

Tandem mass spectrometry profiles showing extracted ion counts (XIC) for isolated fragments from (A) fractionated *E. coli* peptidoglycan and including LC-MS/MS fragmentation for (B, E) disaccharide tetrapeptide, (C, F) disaccharide tripeptide and (D, G) disaccharide dipeptide.

**Fig. S8. Analysis of quiescence stimulation by tripeptide (K-A-A) and pentapeptide (A-E-K-A-A)**

JJ1886, JJ1886 *ΔoppABCDF* and JJ1886 *ΔmppA* ($10^3$) cells were plated onto M9 minimal agar with 0.2% glucose and was challenged with *S. aureus* pentapeptide (L-Ala-D-Glu-L-Lys-D-Ala-D-Ala) (5 µg), Plates were incubated for 24 h. White arrows indicate position of stimulant addition.

Fig. S1

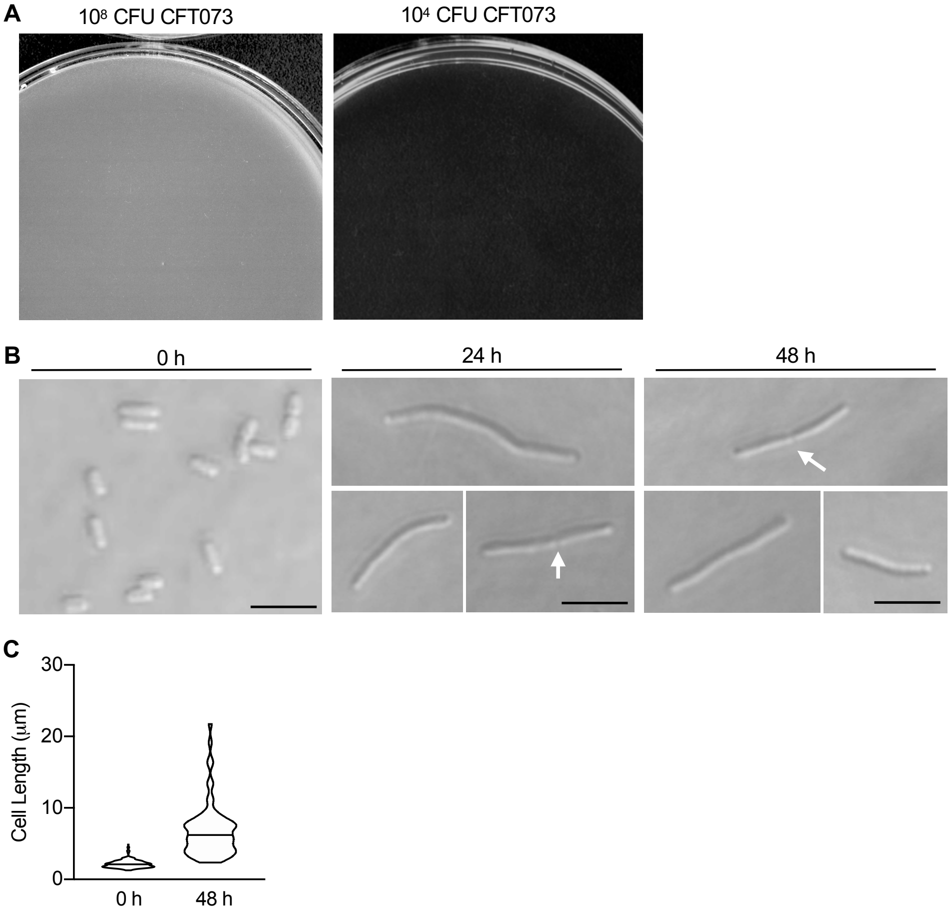
A
$10^8$ CFU CFT073
$10^4$ CFU CFT073
B
0 h
24 h
48 h
C
30
20
10
0
Cell Length (μm)
0 h
48 h

Fig. S2

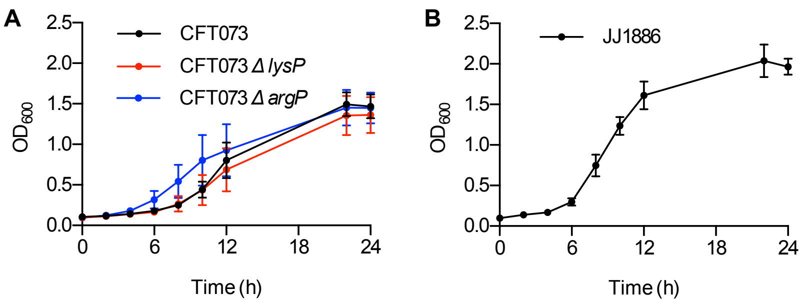
A
CFT073
CFT073 Δ lysP
CFT073 Δ argP
OD600
Time (h)
B
JJ1886
OD600
Time (h)

Fig. S3

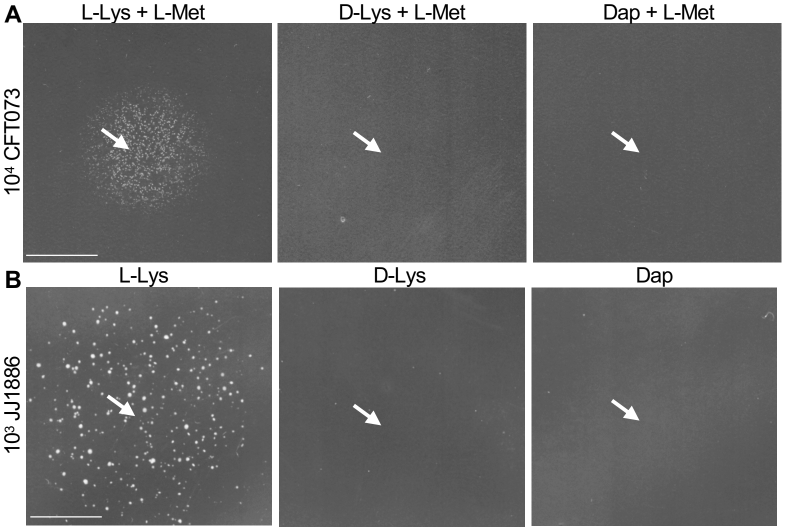
A
L-Lys + L-Met
D-Lys + L-Met
Dap + L-Met
10⁴ CFT073
B
L-Lys
D-Lys
Dap
10³ JJ1886

Fig. S4

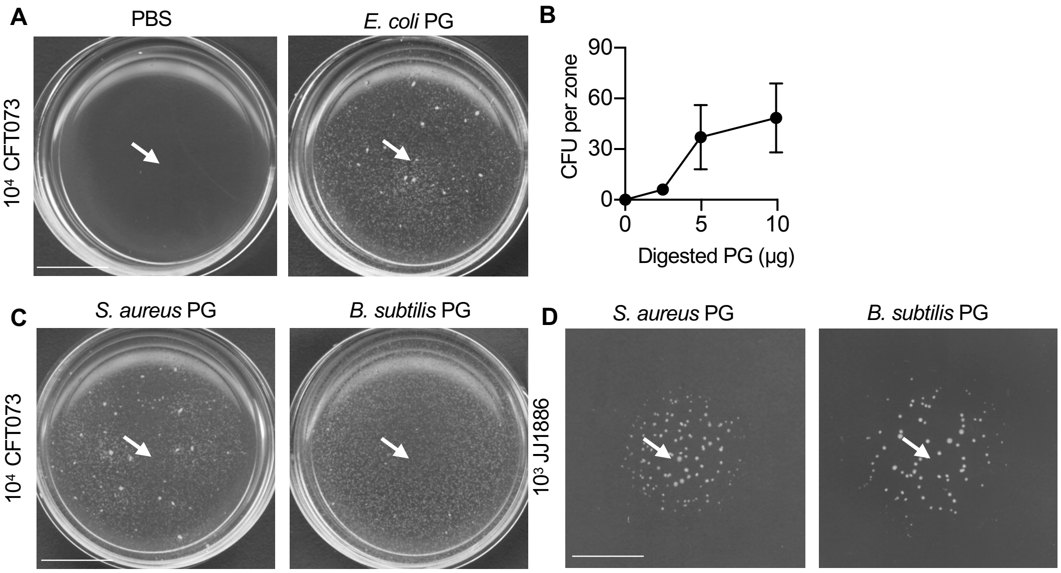
A
PBS
E. coli PG
10⁴ CFT073
B
CFU per zone
90
60
30
0
0
5
10
Digested PG (µg)
C
S. aureus PG
B. subtilis PG
10⁴ CFT073
D
S. aureus PG
B. subtilis PG
10³ JJ1886

Fig. S5

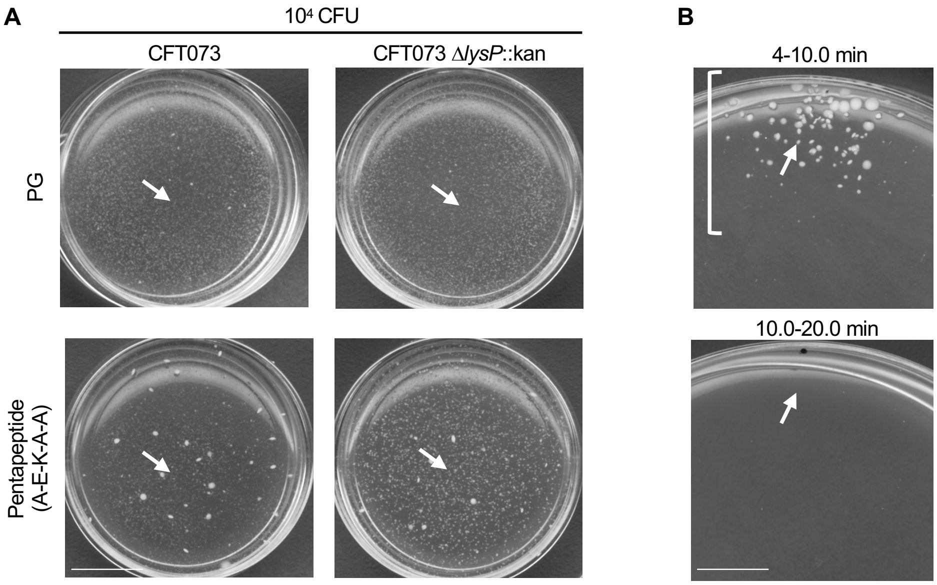
A
10^4 CFU
CFT073
CFT073 ΔlysP::kan
PG
Pentapeptide (A-E-K-A-A)
B
4-10.0 min
10.0-20.0 min

Fig. S6

**A**

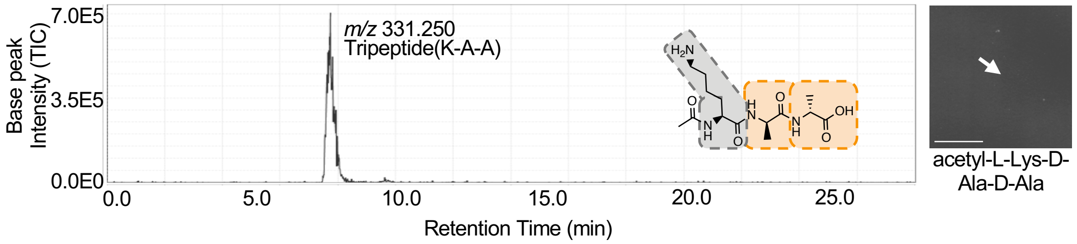
7.0E5
3.5E5
0.0E0
Base peak Intensity (TIC)
0.0
5.0
10.0
15.0
20.0
25.0
Retention Time (min)
*m/z* 331.250
Tripeptide(K-A-A)
acetyl-L-Lys-D-Ala-D-Ala

**C**

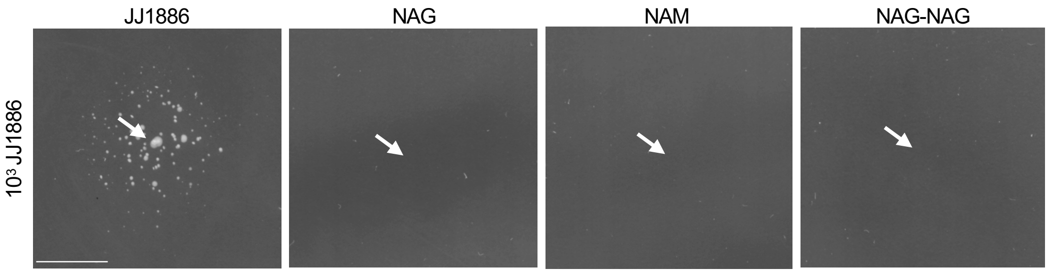
JJ1886
NAG
NAM
NAG-NAG
$10^3$ JJ1886

Fig. S7

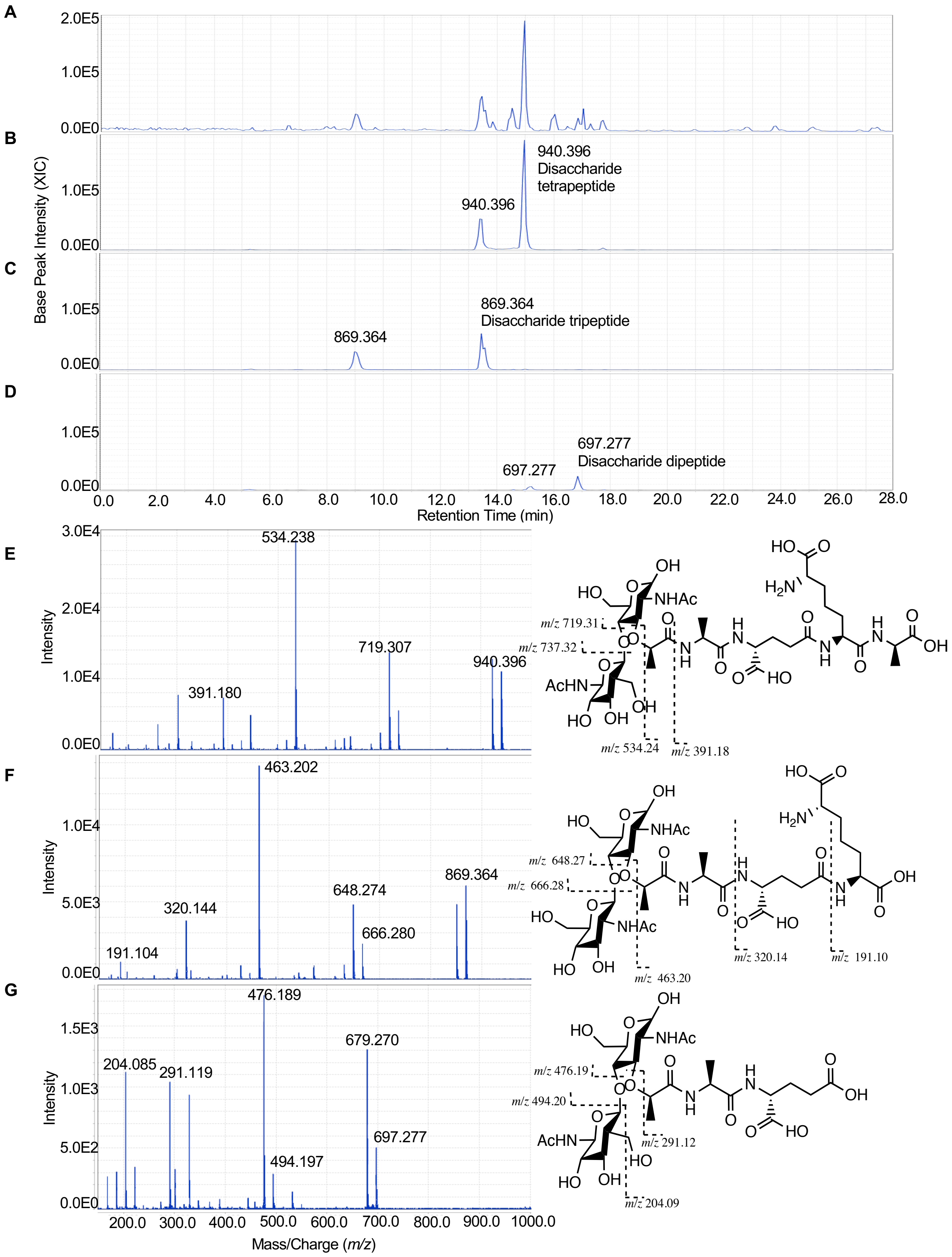
A
B
C
D
E
F
G
Base Peak Intensity (XIC)
940.396
Disaccharide tetrapeptide
869.364
Disaccharide tripeptide
697.277
Disaccharide dipeptide
Retention Time (min)
Intensity
534.238
719.307
940.396
391.180
463.202
648.274
666.280
869.364
320.144
191.104
476.189
679.270
697.277
494.197
204.085
291.119
Mass/Charge (m/z)
m/z 719.31
m/z 737.32
m/z 534.24
m/z 391.18
m/z 648.27
m/z 666.28
m/z 463.20
m/z 320.14
m/z 191.10
m/z 476.19
m/z 494.20
m/z 291.12
m/z 204.09

Fig. S8

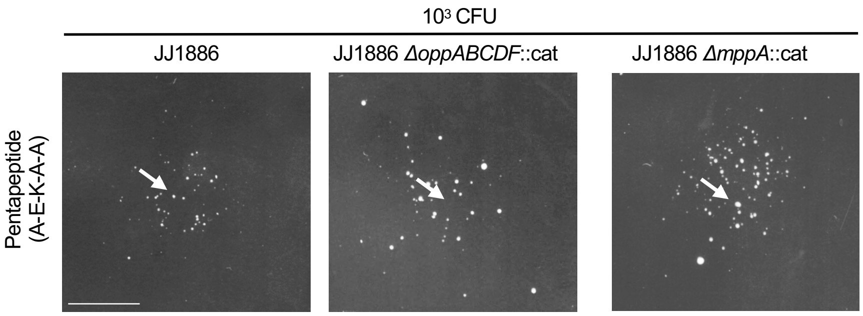
10³ CFU
JJ1886
JJ1886 ΔoppABCDF::cat
JJ1886 ΔmppA::cat
Pentapeptide (A-E-K-A-A)
